# RNA-seq analyses: Benchmarking differential expression analyses tools reveals the effect of higher number of replicates on performance

**DOI:** 10.1101/2020.06.10.144063

**Authors:** Samson Pandam Salifu, Hannah Nyarkoah Nyarko, Albert Doughan, Haward Keteyo Msatsi, Isabel Mensah, Abdul-Rahman Adamu Bukari

**Author notes:** Corresponding author: Samson Pandam Salifu.

## Abstract

The introduction of several differential gene expression analysis tools has made it difficult for researchers to settle on a particular tool for RNA-seq analysis. This coupled with the appropriate determination of biological replicates to give an optimum representation of the study population and make biological sense. To address these challenges, we performed a survey of 8 tools used for differential expression in RNA-seq analysis. We simulated 39 different datasets (from 10 to 200 replicates, at an interval of 5) using compcodeR with a maximum of 100 replicates. Our goal was to determine the effect of varying the number of replicates on the performance (F1-score, recall and precision) of the tools. EBSeq and edgeR-glmRT recorded the highest (0.9385) and lowest (0.6505) average F1-score across all replicates, respectively. We also performed a pairwise comparison of all the tools to determine their concordance with each other in identifying differentially expressed genes. We found the greatest concordance to be between *limma voom treat* and *limma voom ebayes*. Finally, we recommend employing edgeR-glmRT for RNA-seq experiments involving 10-50 replicates and edgeR-glmQLF for studies with 55 to 200 replicates.

**Author summary:** Downstream analysis of RNA-seq data in R often poses several challenges to researchers as it is a daunting task to choose a specific differential expression analysis tool over another. Researchers also find it challenging to determine the number (replicates) of samples to use in order to give comparable and accurate results. In this paper, we surveyed eight differential expression analysis tools using different number of replicates of simulated RNA-seq count data. We measured the performance of each tool and based on the recorded F1-scores, recall and precision, we made the following recommendations; consider edgeR-glmRT and edgeR-glmQLF for replicates of 10-50 and 55-200 respectively.

## Introduction

Since the introduction of RNA sequencing in the mid-2000s, undoubtedly, there has been an exponential increase in RNA-seq data generation with an equivalent rise in the development of algorithms for differential gene expression (DGE) analyses with varying performances. These methods seek to make data analyses relatively easier and address complex biological questions with greater levels of statistical confidence. However, the challenge still remains the selection of optimal DGE tools and sample size calculations for optimal accuracy. This makes the selection of tools and sample sizes for optimum analyses a very crucial but daunting task.

Over the years, several research articles have been published that address the lack of consensus among DGE tools. Examples of these are the works of Seyednasrollah *et al* (1) who performed a systematic comparison of some popular DGE tools and provided recommendations for choosing the optimal tool. Rapaport *et al* (2) assessed a number of tools based on the performance of normalization, false-positive rates and the effect of sequencing depth and sample replication on DGE analyses. Kvam *et al* (3) compared the ability of edgeR, DESeq, baySeq, and TSPM to detect DEGs from both simulated and real RNA-seq data. Germain *et al* (4) assessed the effect of library size on quantification and DEA and went on to create an R package (RNAontheBENCH) and a web platform that could be used for benchmarking RNA-seq quantification and differential expression methods. The influence of the number of replicates, sequencing depth, and balanced versus unbalanced sequencing depth within and between groups using Cufflinks-Cuffdiff2, DESeq and edgeR was explored by Zhang *et al* (5). They concluded that edgeR performed better than DESeq and Cuffdiff2 in terms of its ability to detect true positives and recommended that Cuffdiff2 should not be used if sequencing depth is low (i.e. <10 million reads per individual sample).

Furthermore, most DGE analyses have been limited to designed experimental studies (eg. treated cell lines vs untreated cell lines), which characteristically utilize small (< 12) replicate samples limiting the power of statistical inference.

In this study, we performed a comparative analysis of the performance of eight (8) DGE tools including ABSseq (6), ALDEx2 (7), edgeR (8), limma (9), EBSeq (10), sSeq (11), baySeq (12) and DESeq2 (13). The tools were assessed with a total of fourteen (14) different methods (algorithms used by the tools to identify DEGs) on simulated datasets generated with CompcodeR (14). We also determined the effect of varying the number of replicates per group (sample size) on the performance of each method.

Unique to our study, we used a very high number of samples (20 to 400, at an interval of 5) to assess the performance of the DGE tools and to the best of our knowledge, this is the first study to employ such huge sample sizes to find DEGs in bulk RNA-seq analysis. Our sample sizes were chosen to reflect the current trends of experimental designs for particularly cancer research and population-based studies as smaller numbers of sample replicates are not enough to characterize the high heterogeneity in such studies.

## Results and discussion

The performances of 14 DGE methods of 8 tools for RNA-seq analyses on varying numbers of replicates in two groups were critically assessed.

### Tool Selection

The methods (Limma voom (treat), Limma voom (eBayes), Limma trend (treat), Limma trend (eBayes), edgeR Exact, edgeR likelihood ratio test, edgeR quasi-likelihood F-test, DESeq2, baySeq, EBseq, ALDEx2, sSeq, ABSseq (Classic) and ABSseq (aFold)) were selected based on the following criteria: Firstly, Poisson distribution assumes equal mean and variance across the data and this is atypical of RNA-seq count data which present different means and variance. We based seelction on the above to eliminate all tools that employ Poisson distributions rather than Negative Binomial. Secondly, we selected open-source software packages, which have their source code released under a license that grants users the right to make changes and redistribute the software under certain conditions (15). These software packages are usually robust and have diverse perspectives. We based our choice on this to also eliminate tools that are not open-sourced.

### Comparison of tool performances

All tools were run with default parameters for both DGE and selection of significant genes. The calculated F1-Score for each method was a priority for the performance check as it gave a weighted average of precision and recall taking into account both false negatives and positives. All tools used in this study identified between 265 and 2500 DEGs across all datasets (**Fig 1**).

**Fig 1.**
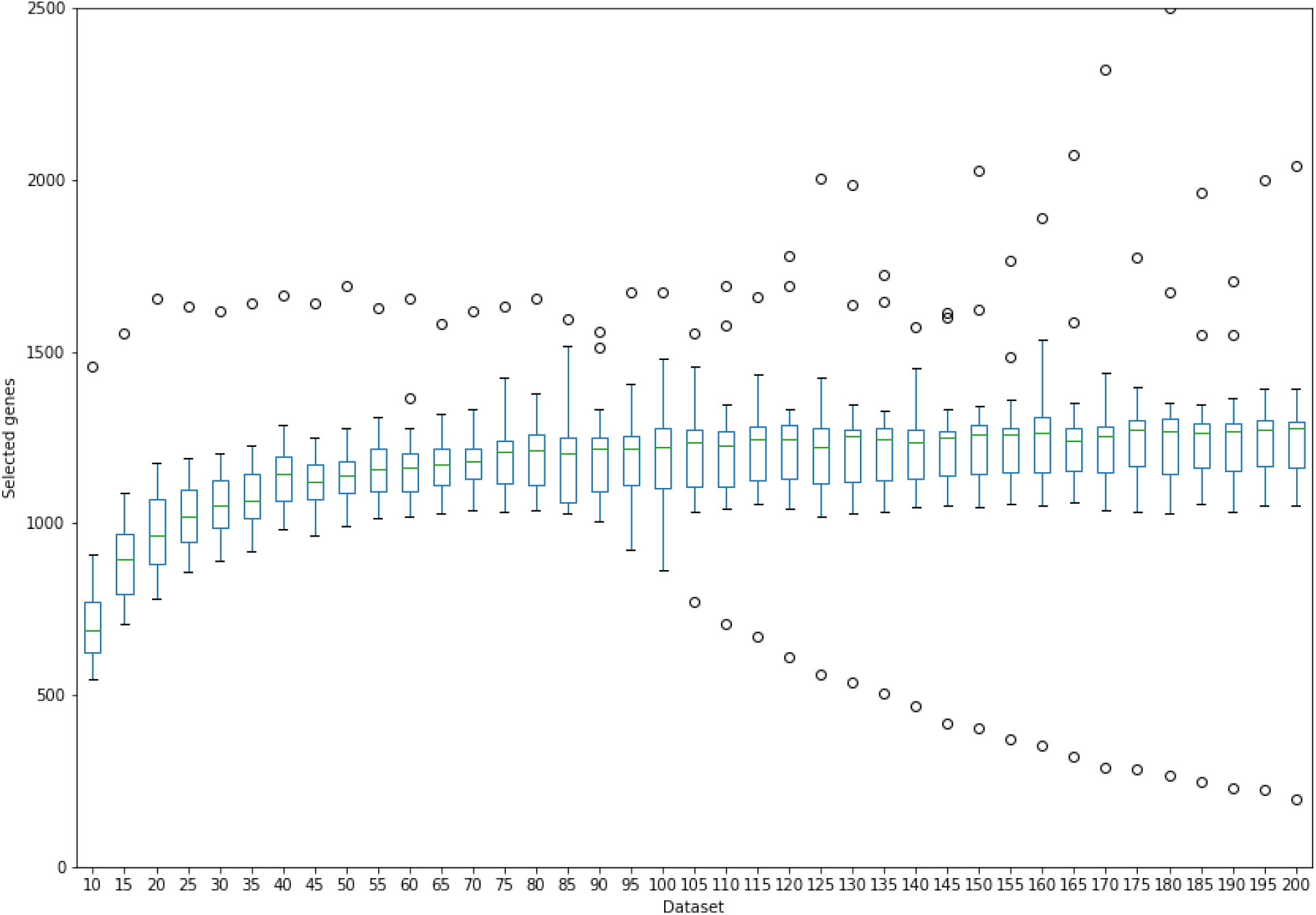
A boxplot comparing the total number of DEGs (y-axis) for all the tools studied at various sample sizes (x-axis). In most of the tools, as the sample sizes increase, the total number of differentially expressed genes also increases steadily. There was an overall trend of increase in the number of DEGs as the sample sizes increase.

EBSeq and edgeR-exact recorded the lowest (266) and the highest (2500) number of DEGs which included the highest number of false negative and positives, respectively.

**Fig 1** presents an overview of the distribution of DEGs identified by all tools at different sample sizes. Irrespective of the tool used, the number of significant genes increases with increasing sample size until a point where it plateaus (after dataset 105). From **Figs 2** and **4**, F1-Scores and recall of EBSeq increased with increasing sample sizes until dataset 75, where it begins to drop, indicating that EBSeq will work best with experimental designs of smaller sample sizes (<75) and might not be appropriate for analyses with larger sample sizes (>75).

**Fig 2.**
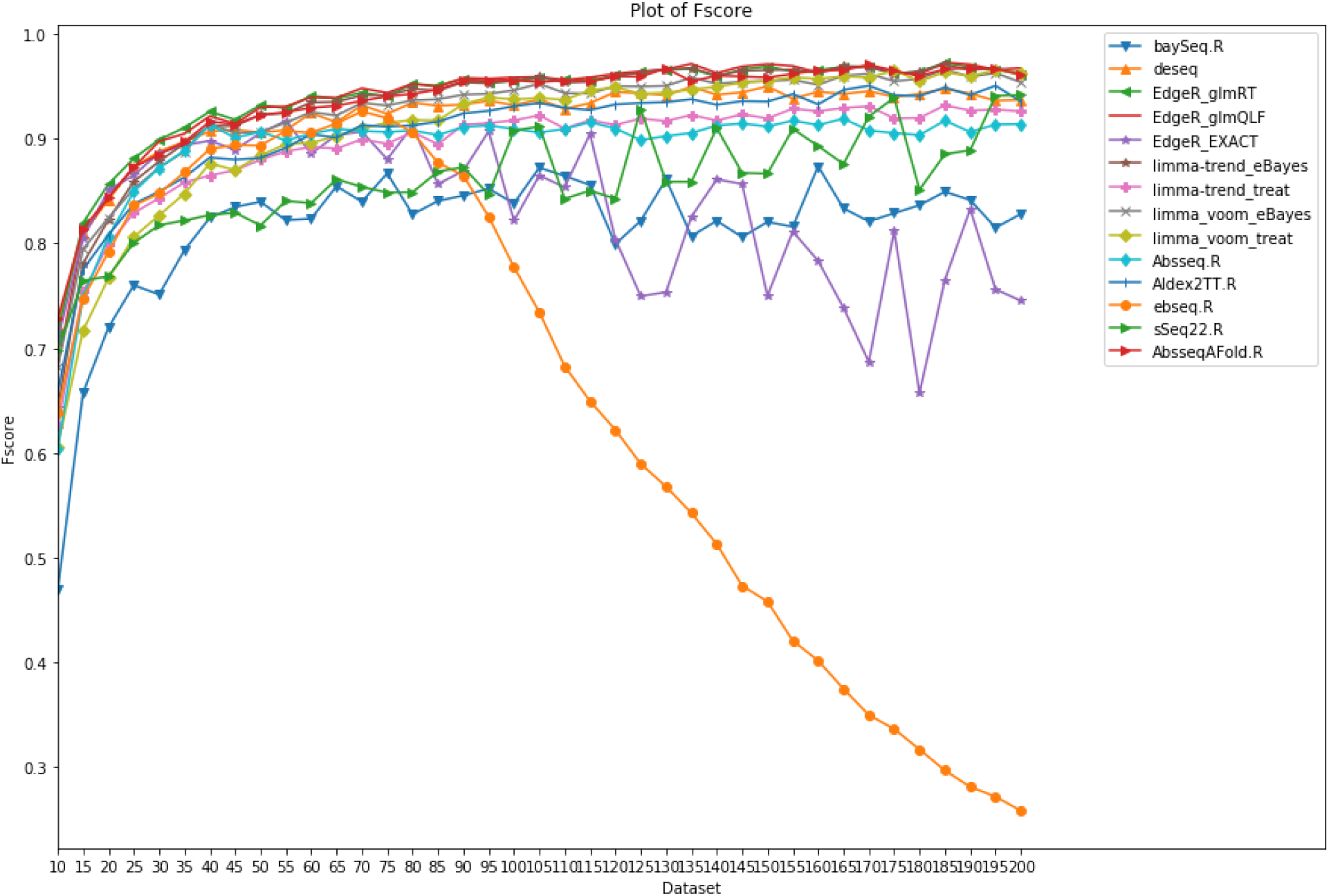
Comparison of F1-scores of all the methods at different sample sizes. The F1-scores of the DGE tools increased with accompanying increase in sample size. The F1-score for EBSeq declines for sample size > 75.

Limma-voom (*treat* and *ebayes*) and limma-trend (ebayes) gave relatively higher F1-scores, compared to limma-trend (*treat)*, which produced poorer F1-scores. This is consistent with those reported by Costa-Silva *et al* (16) (**Fig 2**).

The outstanding performance (F1-scores) of *ebayes* against *treat* method could be attributed to its ability to control the false positive rate (keeping a good balance between sensitivity and precision). The *treat* method on the other hand compensates its poor sensitivity with very high precision scores.

Generally, larger sample sizes give more reliable results with greater precision and power (17). Furthermore, work by Liu *et al* (18) and Busby *et al* (19) established that increasing the number of replicates in an RNA-seq analysis usually leads to more robust results. This is evident as the majority of the tools showed increasing precision and F1-scores with increasing sample sizes (**Fig 2**). Moreover, across the majority of tools assessed (ABSseq, ALDEx2, baySeq, sSeq and Limma), there seemed to be a higher change in performance from sample sizes of 20 to 50 after which the F1-Scores increase steadily until they reached a plateau. This points to the fact that using a sample size of 100 and above per group in DEA might be optimal.

edgeR, DESeq2 and EBSeq recorded a slight decrease in performance at the highest sample size (200), which is in contrast with Biau *et al*’s (17) assumption (larger sample is equivalent to greater precision and power). This could be accounted for by the rise in the false positives and the false negatives recorded at this sample size.

Whilst limma and ABSseq (*aFold*) are amongst the best performing tools overall, this evaluation indicates that for analyses requiring high precision, limma-trend, ABSseq (classic) and edgeR would be the best option whilst sSeq and DESeq2 could be optimal for analyses prioritizing recall (**Fig 4**). This is in line with a study by Lamarre (20) who found that the DESeq2 pipeline seems to prioritize recall while limma prioritizes precision. It is also worth noting that whilst recall of all the tools (except EBseq) seems to be strongly dependent on sample size, precision is independent of sample size.

**Fig 3:**
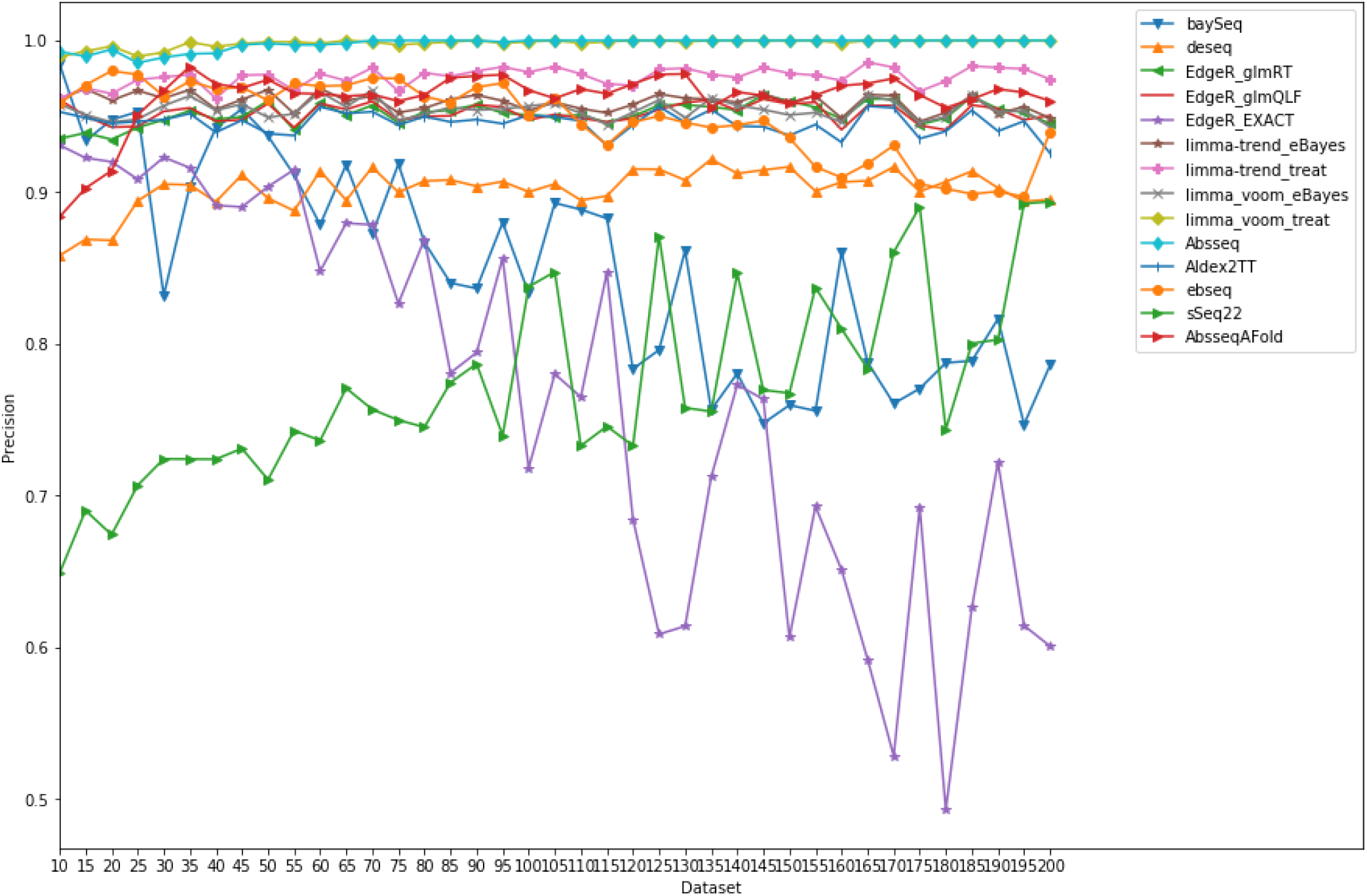
Comparison of precision values for all the methods at different sample sizes. edgeR-exact recorded low values of precision across all the datasets, followed closely by sSeq and baySeq. ABSseq, limma voom (treat) and limma trend (treat), all gave consistently higher precision values.

**Fig 4.**
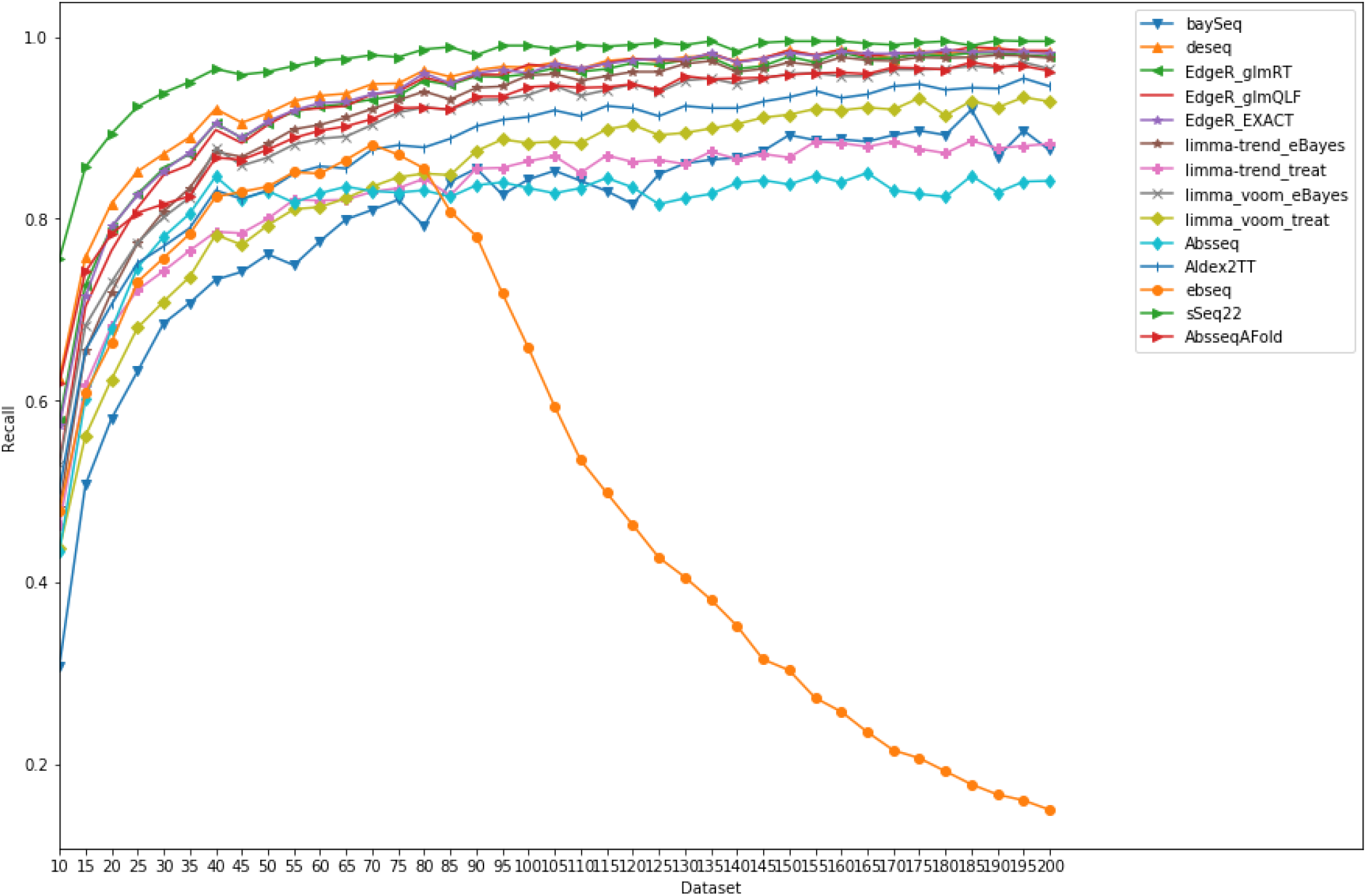
Comparison of recall scores by all the methods at different sample sizes. Overall, sensitivity increased gently across the various datasets in all methods, with sSeq (Dataset_100) recording the highest (0.9872). The least sensitive method was limma voom (treat) (0.4296) at a sample size of 20.

We also investigated the concordance between a pair of DGE tools using a heatmap. This was generated from data at sample size 75, where almost all the tools had an optimal performance and therefore typical of the F1-score of the entire datasets. We plotted a heatmap using the top 1000 most significant DEGs from each tool based on adjusted p-values or FDR. The heatmap was used to identify tools that had the highest number of common genes. We surmise that these tools employ similar approaches in identifying DEGs. limma voom *(treat)* and limma voom *(ebayes)*. identified the highest number (994) of DEGs while edgeR (QLF) and ABSeq (aFold) recorded the least (808) (**Fig 5**).

**Fig 5.**
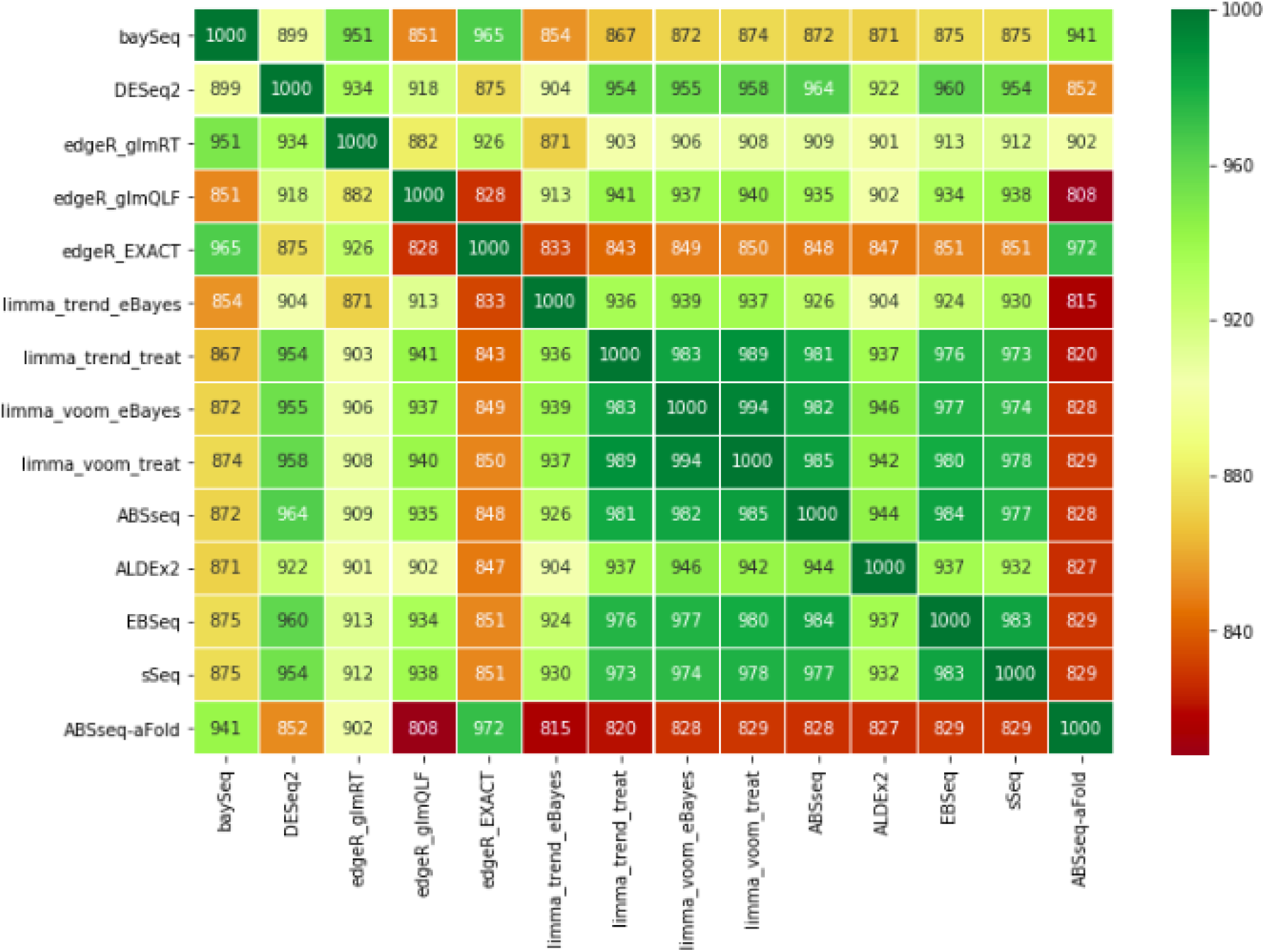
A heatmap showing the pairwise concordance between DGE tools. The highest concordance was between limma voom (treat) and limma voom (ebayes) while the lowest was between edgeR (QLF) and ABSeq (aFold).

*limma voom treat* and *limma voom ebayes*. identified the highest number (994) of DEGs while edgeR (QLF) and ABSeq (aFold) recorded the least numbers (808).

Finally, we compared the averaged F1-scores of all the methods in order to identify the overall best- and worst-performing tools (**Fig 6**). Based on this, the best tool was edgeR-glmRT (avg. F1-score = 0.938498), which would be ideal for DGE analysis of RNA-seq data involving a maximum of 100 replicates.

**Fig 6.**
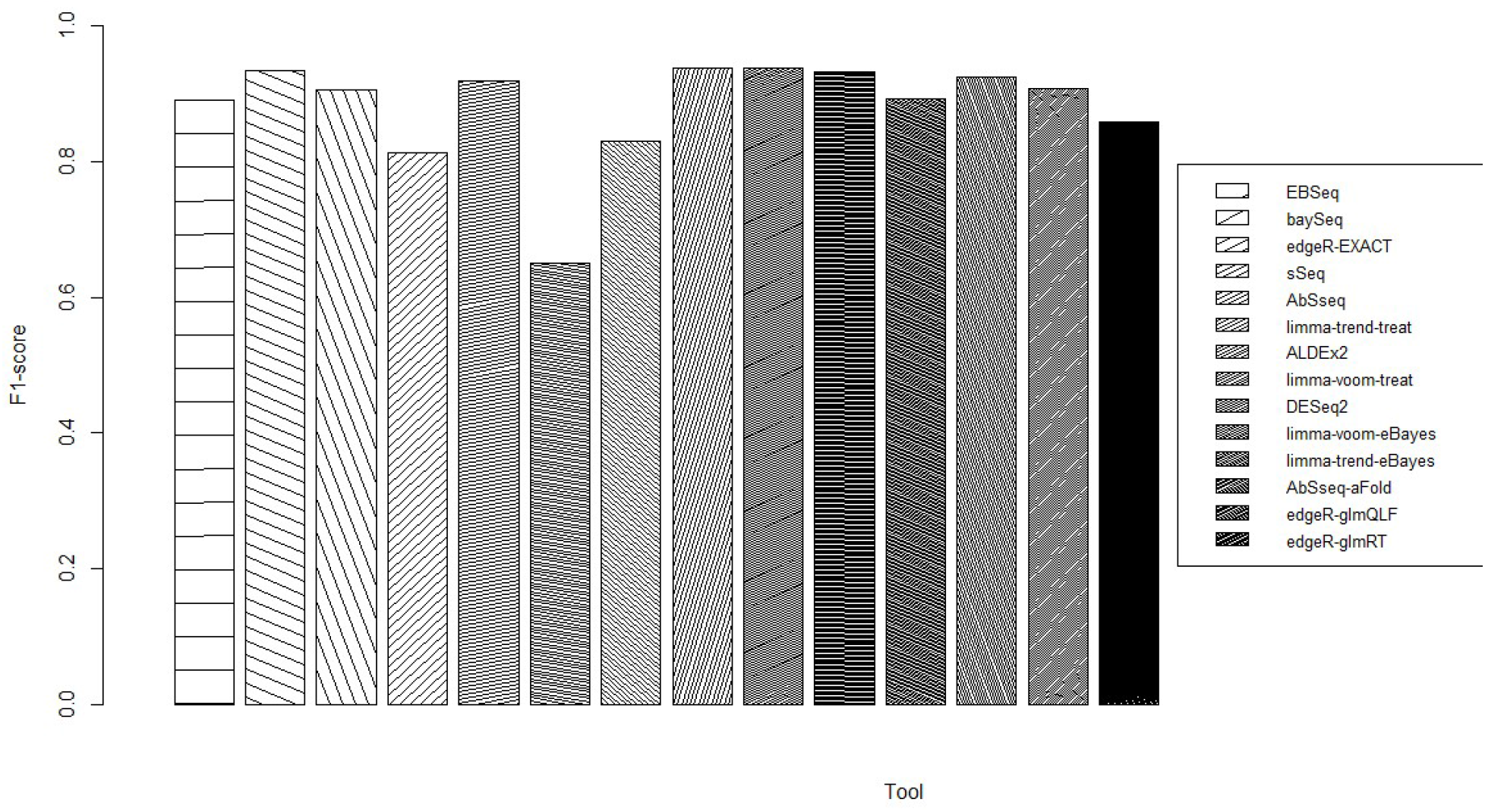
Performance (F1-score) measure of DGE tools. edgeR (glmRT) and EBSeq were the overall best- and worst-performing tools respectively based on average F1-score.

An alternative would be edgeR (glmQLF) ((avg. F1-score = 0.937598)) and AbSseq (aFold) (avg. F1-score = 0.934508). EBSeq (avg. F1-score =0.650498), baySeq (avg. F1-score = 0.813696) and edgeR (EXACT) (avg. F1-score = 0.830104) (**Fig 6**) are not recommended for research studies involving up to 100 replicates.

## Conclusion

We presented a detailed review of different methods used for differential expression analysis of simulated count data from RNA-seq experiments. After evaluating the effect of increasing the number of replicates on the performance (F1-score, recall and precision) of DGE tools, we made the following recommendations that may be relevant to researchers employing diverse replicates in their RNA-seq experiments; for replicate numbers between 10 and 50, edgeR-glmRT produces better results while edgeR-glmQLF was better for replicates between 55 and 200. We did not identify among the evaluated methods a tool that produces optimum results in all the performance measures for the evaluated replicate numbers.

For recall, precision and F1-score, sSeq (0.970913), limma_voom_treat (0.998309) and sSeq (0.970913) produced the best results, respectively. Depending on the objectives of a particular research, investigators can prioritize one performance measure over the other and employ the appropriate recommended method.

## Materials and Methods

### Tool selection criteria

Several tools have been developed for DGE in RNA-seq analyses. These tools adopt diverse distributions and methods to identify DEGs. Popular among these are the Negative Binomial (NB) and Poisson distributions. Upon surveying over 35 tools, 8 were selected based on our set criteria: tools that follow a Negative Binomial Distribution and are open-source. Tools that employ Poisson distribution were not considered since it assumes equal variance across a given dataset, which is atypical of RNA-seq data.

### Data Simulation and datasets

We simulated an RNA-seq count data set (dataframe containing the number of reads mapping to each genomic feature of interest in each of the samples) using compcodeR’s *generateSyntheticData* function (14), with the mean values sampled from values estimated from the Pickrell (21) and Cheung (22) data sets. Gene counts were simulated following the Negative Binomial distribution.

The simulated datasets contained 12,500 gene counts with two groups of 10 replicates each, where 10% of the simulated genes were differentially expressed between the two groups. Counts were also simulated with the same dispersion in the two groups, and no outlier counts were introduced. The datasets were filtered to exclude all genes with total counts of 0 and the DEGs were equally distributed between up regulated and down regulated genes. The above simulation approach was repeated for replicates of 15 to 200 (with an interval of 5).

A set of randomly generated gene names obtained from biomaRt (23) was appended as row names to the count table of the dataset. This newly generated count matrix was further used for DGE analyses. From the dataset generated, the truth set was extracted from the differential.expression column of the variable.annotations table, where “1” and “0” represented differentially expressed and non-differentially genes, respectively. The group designation was also extracted from the sample.annotations table, which contained the group for each sample.

### Differential Gene Expression Analysis

A brief description of the 8 different tools is presented below:

#### DESeq2

A DESeqDataSet object was created from the matrix of counts and metadata using DESeqDataSetFromMatrix function for counts data. The DESeq function was then run on the object created to perform DGE analysis. This was followed by building the results table using the results function. MA plots were then generated from the results obtained. Finally, the selection of DEGs was performed and genes with adjusted p-values of less than 0.1 (default for DESeq2) were considered to be significant and differentially expressed.

#### limma

Firstly, a DGEList object was created using edgeR package in R. Trimmed mean of M values (TMM) normalization method was used on the counts data since it performs well in comparative studies (24). Counts were then converted to logCPM values using edgeR’s cpm function. The logCPM values were then used in the entire analyses using limma-trend pipeline with eBayes and treat methods. The voom transformation was applied to the normalized DGEList object to create an Elist object, which was finally used in eBayes and treat methods for DGE analyses. Genes with adjusted p-values of less than 0.05 (default) were selected as significantly differentially expressed.

#### edgeR

A DGEList object was created from the matrix of counts, ensued by normalization using TMM. Prior to DGE analyses with classic edgeR approach (exact), quasi-likelihood F-test and the likelihood ratio test, the dispersions were estimated. Genes with an adjusted p-value of less than 0.05 were selected as significantly differentially expressed.

#### baySeq

DEA commenced with the creation of a countData object from the simulated data and the already defined groups and replicates. We then inferred the library sizes. Prior to obtaining posterior probabilities and estimating proportions of differentially expressed counts, we estimated prior probabilities on countData object using the negative binomial model. Genes with an adjusted p-value of less than 0.05 (default) were selected as significantly differentially expressed.

#### EBSeq

Working with EBSeq requires an estimation of library sizes for each sample. Here, the library sizes were obtained by a median normalization function. The data was converted into a matrix with all the genes explicitly stated as the row names. The function EBTest was used, which considers the count data, the conditions, the library sizes and the expected number of iterations. GetPPMat and GetDEResults were used to output the results and to extract the significantly differentially expressed genes (p-value less than or equal to 0.5).

#### ALDEx2

It estimates per-feature technical variation within each sample using Monte-Carlo instances drawn from the Dirichlet distribution. During the analyses, we first set the comparison groups, which is a vector of conditions in the same order as samples in the counts dataset. We then performed a t-test and used 128 Monte-Carlo instances as recommended by Gloor (7) for t-test analyses. DEGs were extracted by setting a threshold of adjusted p-value to less than 0.1 (default).

#### sSeq

The function nbTestSH was used to obtain the regularized dispersion estimates and perform the exact tests. P-values were corrected with the Benjamini-Hochberg method using the p.adjust function. Significantly differentially expressed genes were extracted by setting a threshold of adjusted p-value less than 0.05 (default).

#### ABSseq

We created an object by providing a count matrix (from the simulated gene counts) and the defined groups. The ABSDataSet function includes a parameter for normalization, which by default is qtotal. To identify differentially expressed genes, the ABSseq function was used. This ran a default analysis by calling all required functions in the background. In an alternative approach (aFold), DEGs were called via log fold-change. It uses a polynomial function to model the uncertainty (variance) of read count, and thus takes into consideration the variance of expression levels across treatments and genes. In this approach, useaFold was set to ‘TRUE’. In both approaches, differentially expressed genes were extracted by setting a threshold of adjusted p-value < 0.05 (default).

### Performance metrics measurement

The simulated datasets and ‘truthset’ were used to assess the performance of each method based on precision, recall and F1-score. We also explored the effect of varying the number of replicates in both groups on the performance of each method. Finally, we performed a pairwise comparison on the number of DEGs recorded for each tool.

## Acknowledgements

The authors would like to acknowledge Richard Ansah and Peter Amoako-Yirenkye for their dedication and maintenance of our servers and Nicola J. Mulder for language revision and suggestions

## Declarations

### Ethics Approval and consent to participate

Not applicable

### Consent for publication

Not applicable

### Availability of data and materials

Supplementary information can be obtained from https://github.com/h3aknust/Assessing-differential-expression-analyses-tools

### Authors’ contributions

Conceptualization, SPS; Methodology, SPS, AD, HNN, IM, HKM and AAB; Validation, SPS; Formal Analysis and Data Curation, AD, HNN, IM, HKM and AAB; Original Draft Preparation, AD, HNN, IM, HKM and AAB, Review and Editing, SPS, AD, HNN, IM, HKM and AAB; Resources, Project Administration and Funding Acquisition, SPS.

